# Cardiac-Detargeted MyoAAV Enables Systemic Nrl-Mediated Fast Myofiber Remodeling and Hypertrophy Across Multiple Skeletal Muscles

**DOI:** 10.64898/2026.04.20.719776

**Authors:** Keisuke Hitachi, Shunya Sadaki, Masato Watanabe, Ryosuke Tsuji, Atsushi Kubo, Yuki Yamasaki, Yuri Kiyofuji, Masafumi Inui, Takashi Kudo, Tomohiko Suzuki, Satoru Takahashi, Kunihiro Tsuchida, Ryo Fujita

## Abstract

Inducing fast myofiber programs offers therapeutic potential for skeletal muscle disorders such as sarcopenia, where fast myofibers are preferentially lost. Engineered muscle-specific AAV (MyoAAV) vectors enable efficient transduction of skeletal muscles after systemic administration; however, cardiac transgene expression limits applications requiring skeletal muscle-selective delivery. We generated modified MyoAAV vectors by incorporating cardiac-specific miR-208a target sequences into the transgene 3′UTR. This design markedly suppressed cardiac expression while preserving skeletal muscle output, with target-site variation enabling tunable trade-offs between cardiac detargeting and skeletal muscle expression levels. We validated this platform using neural retina leucine zipper (Nrl), a large Maf transcription factor regulating type IIb myofiber identity. Systemic delivery of conventional MyoAAV-Nrl caused severe cardiac hypertrophy and uniform lethality within one month. Conversely, incorporating miR-208a target sequences prevented detectable hypertrophy and eliminated mortality during the experimental observation period. This modification significantly reduced cardiac *Nrl* expression while maintaining skeletal muscle levels, successfully promoting type IIb myofiber formation and hypertrophy across multiple skeletal muscles. These findings demonstrate that miR-208a-mediated cardiac detargeting combined with MyoAAV-Nrl enables safe systemic induction of fast myofiber remodeling and hypertrophy, establishing a platform for gene therapies targeting skeletal muscle disorders associated with fast myofiber loss.

## Introduction

Skeletal muscle is the most abundant tissue in the body and plays critical roles in locomotion, posture, thermogenesis, and metabolic homeostasis. In addition to these mechanical and metabolic functions, skeletal muscle communicates with other organs through circulating factors such as myokines and thereby serves as a major determinant of systemic health. However, muscle wasting frequently occurs under catabolic conditions, including sepsis, cancer cachexia, and aging.^1,2^

Skeletal muscle is primarily composed of multinucleated myofibers that exhibit remarkable heterogeneity in contractile and metabolic properties. Adult myofibers are broadly classified into fatigue-resistant slow-twitch (type I) fibers and rapidly fatiguing fast-twitch (type II) fibers, most commonly based on their expression of myosin heavy chain (MyHC) isoforms.^3^ Slow type I myofibers are characterized by high mitochondria content and oxidative metabolism and predominantly express MyHC I, encoded by *Myh7*. Fast type II myofibers comprise IIa, IIx, and IIb subtypes, defined by the expression of MyHC isoforms (MyHC IIa, IIx, and IIb) encoded by *Myh2*, *Myh1*, *Myh4*, respectively. These subtypes display progressively faster contractile properties and greater glycolytic capacity. In conditions associated with impaired motor innervation, such as aging, Duchenne muscular dystrophy (DMD), amyotrophic lateral sclerosis (ALS), fast myofibers are particularly vulnerable, and rodent models typically exhibit marked atrophy and selective loss of type IIb myofibers.^4–9^ These observations highlight the importance of developing therapeutic strategies that efficiently target fast myofibers and preserve their identity and function in muscle wasting diseases.

Recombinant adeno-associated virus (AAV) vectors have emerged as a promising platform for skeletal muscle gene therapy because they enable efficient and durable in vivo transduction.^10–12^ However, conventional AAV serotypes often lack sufficient skeletal muscle specificity after systemic delivery potentially mediating substantial off-target transduction in non-target tissues. To overcome this limitation, engineered muscle-tropic capsids such as MyoAAV have been developed to enhance gene delivery to striated muscles, including skeletal and cardiac muscle, in disorders such as DMD.^13^ Despite this advance, residual cardiac tropism remains an important limitation of MyoAAV, particularly in applications requiring selective gene delivery to the skeletal muscle. Off-target cardiac expression may not only reduce therapeutic specificity but also raise safety concerns when the transgene exerts deleterious effects in the heart. Therefore, further optimization of MyoAAV is required to achieve greater skeletal muscle specificity with reduced cardiac transduction, thereby improving its utility for preserving muscle function in muscle diseases and aging.

However, vector optimization alone is insufficient; therapeutic success additionally depends on the choice of transgene. We previously identified large Maf transcription factors as essential regulators of fast type IIb myofiber identity in mice.^14^ Large Maf family proteins (Mafa, Mafb, Maf, and Nrl) are basic region–leucine zipper (bZIP) transcription factors, in which the basic region and leucine zipper mediate DNA binding and dimerization, respectively, while N-terminal acidic/transactivation domains modulate target gene transcription.^15,16^ These factors recognize Maf recognition elements (MAREs)-extended DNA motifs related to TRE/CRE cores but with characteristic flanking sequence requirements—and can also act through MARE half-sites depending on local sequence context. Among the four large Maf transcription factors found in mammals (Mafa, Mafb, Maf, and Nrl), Mafa, Mafb, and Maf are endogenously expressed in skeletal muscle in both mice and humans, whereas Nrl is not. Nrl is known as a retina-enriched large Maf factor that functions as a key determinant of rod photoreceptor fate during mammalian eye development.^17–19^ While Nrl possesses a distinct domain architecture, lacking the characteristic glycine- and histidine-rich low-complexity region found in Mafa, Mafb, and Maf,^20^ it nevertheless retains the ability to recognize MARE/MARE-like motifs and activate downstream transcription through these cis-elements.^21,22^

In skeletal muscle, muscle-specific ablation of Mafa, Mafb, and Maf leads to an almost complete loss of type IIb myofibers, whereas overexpression of each factor is sufficient to induce type IIb myofibers in the slow muscle.^14^ Mechanistically, each large Maf directly activates *Myh4* transcription through a MARE in its promoter. In addition, recent work has implicated Maf in suppressing atrophy-related genes in denervation-induced muscle atrophy and identified Maf as an important mediator linking motoneuron firing to muscle gene regulation, fiber integrity, and neuromuscular junction (NMJ) maintenance in fast glycolytic fibers.^23,24^ Importantly, large Maf factors are also functionally relevant in human skeletal muscle, as forced expression of MAFA, MAFB, or MAF strongly induces *MYH4* expression in human myotubes.^25^ Together, these findings identify large Maf family as attractive therapeutic cargo for preserving fast-myofiber identity. Although Nrl is not detectably expressed in skeletal muscle under physiological conditions and its role in myofiber-type regulation remains unknown, it retains the ability to recognize Maf-responsive cis-elements. Thus, we hypothesized that Nrl could activate the fast-myofiber gene program and serve as an additional candidate transgene for muscle-directed gene therapy.

In this study, we implemented a MyoAAV-based expression system incorporating tandem miR-208a target sites^26,27^ to reduce cardiac off-target expression after systemic delivery. Using Nrl as a proof-of-concept cargo, we showed that systemic administration of cardiac-detargeted MyoAAV-Nrl enabled widespread induction of type IIb myofibers in both forelimb and hindlimb skeletal muscles. Moreover, this system promoted myofiber hypertrophy, as evidenced by increased myofiber cross-sectional area (CSA), as well as increased body weight. Together, these findings demonstrate that combining miR-208a-mediated cardiac detargeting with MyoAAV-Nrl enables fast myofiber remodeling and hypertrophy across multiple skeletal muscles after systemic delivery.

## Results

### MyomiR target sites enable cardiac detargeting of MyoAAV transgene expression

Systemic MyoAAV delivery can drive robust transgene expression not only in skeletal muscle but also in the heart, complicating skeletal muscle-focused studies in which unintended cardiac expression may confound phenotypic interpretation and safety assessment.^28^ To address this limitation, we incorporated tandem target sites for miR-208b-3p, miR-208a-3p, and miR-499-5p into the 3′ untranslated region (3′UTR) of an MHCK7 promoter-driven EGFP cassette (Fig. 1A), based on prior studies defining these miRNAs as a *Myh* gene-encoded myomiR family associated with cardiac and striated muscle gene regulation.^29^ We administered conventional MyoAAV-EGFP and MyoAAV-EGFP-miRNAs (+208a/b/499) vectors intravenously to ICR mice and harvested tissues one month later for analysis. Gross fluorescence imaging revealed robust EGFP signals in the heart of control mice injected with MyoAAV-EGFP, whereas EGFP fluorescence was markedly reduced in the miRNAs group (Fig. 1B). However, gastrocnemius fluorescence was also reduced in the miRNAs group compared with control (Fig. 1C), suggesting that the myomiR target cassette induces a broader attenuation of transgene expression across both cardiac and skeletal muscle.

**Figure 1.**
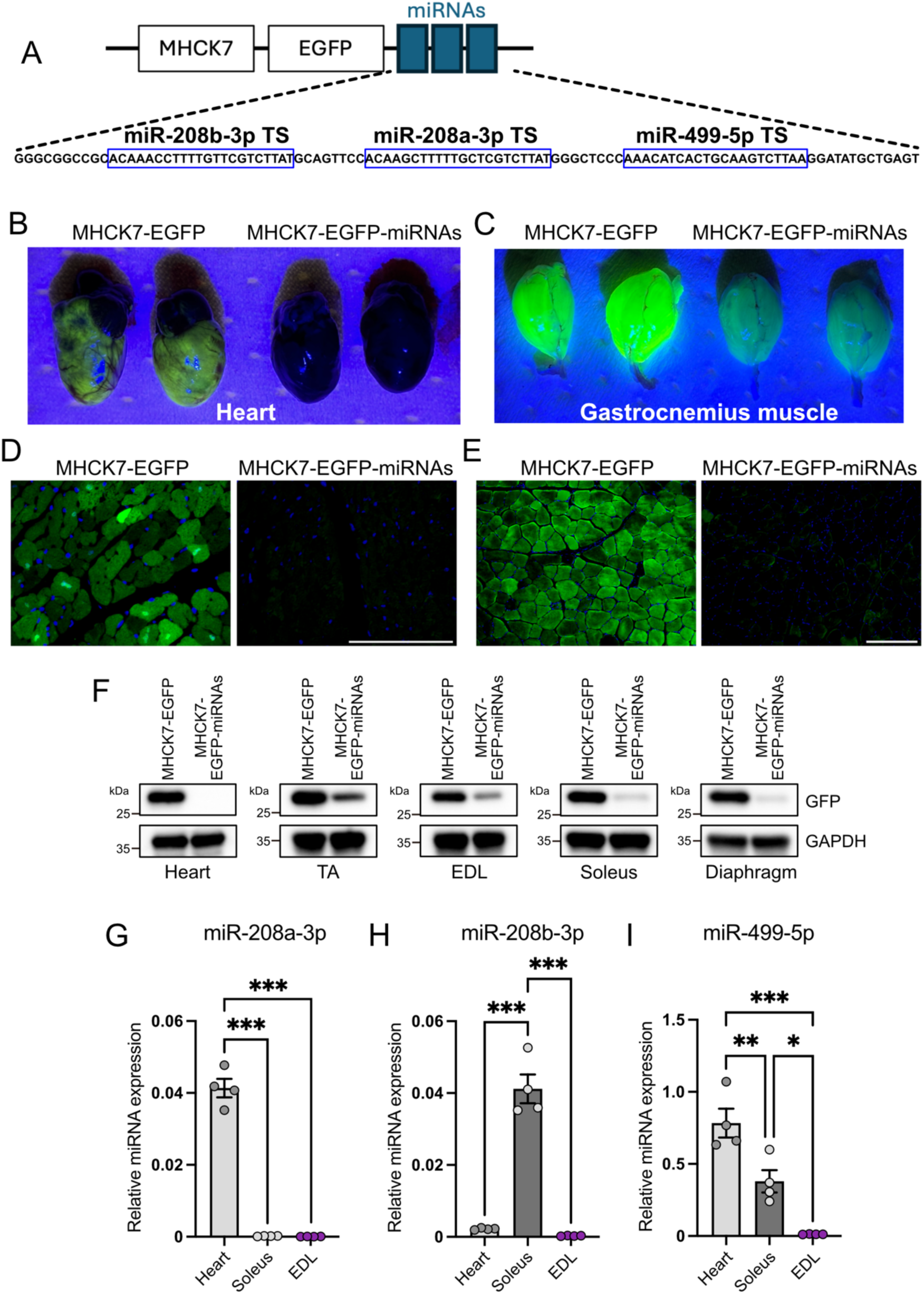
MyomiR target sequences in the 3′UTR suppress cardiac EGFP expression following systemic administration of MyoAAV. (A) Schematic of the MHCK7 promoter-driven EGFP cassette packaged in MyoAAV. Tandem target sites (TS) for miR-208b-3p, miR-208a-3p, and miR-499-5p were inserted into the 3′UTR. Target sequences and spacer sequences are shown. (B) Representative gross fluorescence images of hearts from ICR mice injected intravenously with MyoAAV-EGFP (control) or MyoAAV-EGFP-miRNAs (+208a/b/499) vectors. (C) Representative gross fluorescence images of gastrocnemius muscles. (D) Representative EGFP immunostaining of heart sections from control and miRNAs groups. Nuclei were counterstained with DAPI. Scale bar, 100 µm. (E) Representative EGFP immunostaining of gastrocnemius muscle sections from control and miRNAs groups. Nuclei were counterstained with DAPI. Scale bar, 200 µm. (F) Immunoblot analysis of EGFP protein in heart and indicated skeletal muscles (TA, EDL, soleus, and diaphragm) from control and miRNAs groups. n = 4 mice per group. (G–I) RT-qPCR quantification of mature miR-208a-3p (G), miR-208b-3p (H), and miR-499-5p (I) in heart, soleus, and EDL. n = 4 mice per group. P values were calculated using one-way ANOVA with Tukey’s multiple comparisons test; *P < 0.05, **P < 0.01, ***P < 0.001.

We next examined tissue sections by EGFP immunostaining. Consistent with the gross imaging, EGFP signal was strongly diminished in heart sections from the miRNAs group (Fig. 1D). EGFP immunostaining of gastrocnemius sections also showed reduced signal intensity relative to control (Fig. 1E). To further validate these observations, we assessed EGFP protein levels by immunoblotting. EGFP protein was nearly undetectable in the heart in the miRNAs group, whereas EGFP remained detectable in skeletal muscles including tibialis anterior (TA), extensor digitorum longus (EDL), soleus, and diaphragm, albeit at reduced levels compared with control (Fig. 1F). These results indicate that incorporation of 208a/b/499 target sites enables efficient suppression of cardiac transgene expression, with marked attenuation in skeletal muscle.

To explore whether endogenous myomiR abundance could account for the reduced skeletal muscle expression, we quantified mature miR-208a-3p, miR-208b-3p, and miR-499-5p by RT-qPCR in heart, soleus, and EDL. miR-208a-3p was abundant in the heart but was low in soleus and EDL, whereas miR-208b-3p was detectable in the heart and enriched in soleus. miR-499-5p was robust in the heart, present at approximately half that level in soleus, and was detected at low levels in EDL (Fig. 1G–I). Together, these expression patterns suggest that miR-208b-3p and miR-499-5p activity in skeletal muscle contributes to the reduction of EGFP expression observed in the miRNAs group.

### Optimization of cardiac detargeting using miR-208a target-site repeats

To improve cardiac repression while preserving skeletal muscle expression, we next simplified the miRNA target cassette based on the tissue distribution. Because miR-208a-3p showed the strongest heart enrichment among the tested myomiRs, we replaced the original multi-miRNA cassette with tandem miR-208a-3p target sites. We generated vectors carrying either three target-site repeats (MyoAAV-EGFP-208a×3) or a single target-site repeat (MyoAAV-EGFP-208a×1) in the 3′UTR of an MHCK7 promoter-driven EGFP cassette (Fig. 2A). Following systemic intravenous administration, gross fluorescence imaging showed robust EGFP signals in control animals, whereas EGFP fluorescence in the heart was markedly reduced in both 208a×3 and 208a×1 groups (Fig. 2B). In the gastrocnemius muscles, EGFP fluorescence was comparable between control and 208a×1, whereas the 208a×3 group showed a weaker but still detectable signal (Fig. 2B).

**Figure 2.**
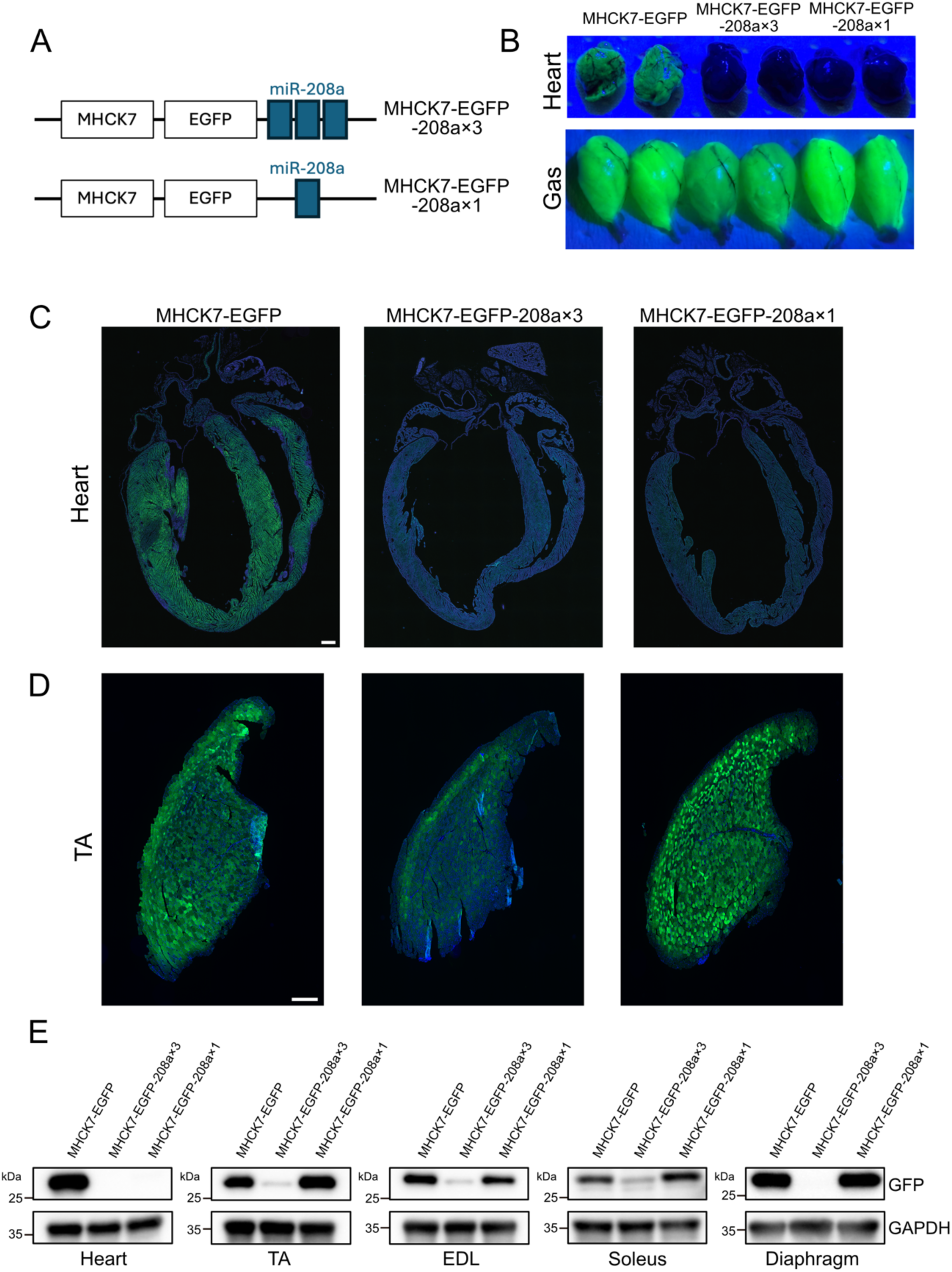
miR-208a target-site repeat number modulates the balance between cardiac repression and skeletal muscle preservation after systemic MyoAAV delivery. (A) Schematic of the MHCK7 promoter-driven EGFP cassette packaged in MyoAAV carrying either three miR-208a-3p target-site repeats (208a×3) or a single miR-208a-3p target-site repeat (208a×1) in the 3′UTR. (B) Representative gross fluorescence images of the heart and gastrocnemius muscles from ICR mice injected intravenously with control, 208a×3, or 208a×1 AAV vectors. (C) Representative cross-sectional images of hearts from control, 208a×3, and 208a×1 groups. EGFP signal was assessed by immunostaining. Nuclei were counterstained with DAPI. Scale bar, 500 µm. (D) Representative TA transverse sections from control, 208a×3, and 208a×1 groups stained for EGFP. Nuclei were counterstained with DAPI. Scale bar, 500 µm. (E) Immunoblot analysis of EGFP protein in the heart and indicated skeletal muscles (TA, EDL, soleus, diaphragm) from control, 208a×3, and 208a×1 groups. n = 4 mice per group.

Consistently, cross-sectional imaging and EGFP immunostaining of heart sections confirmed strong suppression of EGFP in the 208a×3 group, with slightly higher residual signal in the 208a×1 group relative to 208a×3 (Fig. 2C). In contrast, EGFP immunostaining of TA sections indicated that skeletal muscle expression was attenuated by the 208a×3 configuration, whereas the 208a×1 configuration largely preserved EGFP signal in TA (Fig. 2D). We next validated these observations by immunoblotting for EGFP in heart and multiple skeletal muscles (TA, EDL, soleus, and diaphragm). EGFP protein was nearly undetectable in the heart in both 208a×3 and 208a×1 groups, although the 208a×1 group showed a slightly higher residual signal than 208a×3 (Fig. 2E). Consistent with this, high-exposure imaging of the same immunoblot revealed a faint residual EGFP signal in the heart in the 208a×1 group (Supplementary Fig. 1A). However, similar to the multi-miRNA cassette, the 208a×3 configuration also reduced EGFP protein in skeletal muscles, with only faint residual signals detectable. By contrast, the 208a×1 configuration maintained skeletal muscle EGFP levels close to control (Fig. 2E), demonstrating that the repeat number of miR-208a-3p target-site enables a tunable balance between cardiac repression and skeletal muscle expression, such that 208a×3 achieves more stringent cardiac suppression, whereas 208a×1 better preserves skeletal muscle expression. Cre-based reporter analysis revealed cardiac recombination despite the incorporation of miR-208a target-site repeats (Supplementary Fig. 1B, C), indicating that the level of stringency required for cardiac detargeting is highly dependent on the sensitivity of readout and nature of the transgene. These results indicate that, although miR-208a target-site repeats provide effective cardiac repression in protein-based readouts, their performance may be less stringent in recombinase-based applications.

### Cardiac detargeting is robust across purification workflows and adult dosing routes

To assess the practical flexibility of our AAV vector, we compared vector preparation and dosing conditions that could broaden its use in routine in vivo experiments, including adult intraperitoneal (IP) delivery as a practical, less conventional alternative to intravenous (IV) injection. Using the MyoAAV-EGFP-208a×1 vector, we prepared three dosing groups that differed only in vector purification and injection route: Group 1, ultracentrifugation (UC)-purified vector administered intravenously; Group 2, kit-purified vector without UC administered intravenously; and Group 3, UC-purified vector administered intraperitoneally (Fig. 3A). We first assessed transgene expression by immunoblotting for EGFP in the heart and TA. EGFP protein was not detectable in the heart in any group, indicating that cardiac detargeting by the 208a×1 cassette was maintained across purification and delivery conditions (Fig. 3B). In TA, EGFP protein was detected in all groups; however, the EGFP signal in Group 3 (UC purified, IP) was relatively lower than that in Group 1 (UC purified, IV). Densitometric quantification confirmed that EGFP levels in Group 3 were approximately one third of those in Group 1 (Fig. 3C).

**Figure 3.**
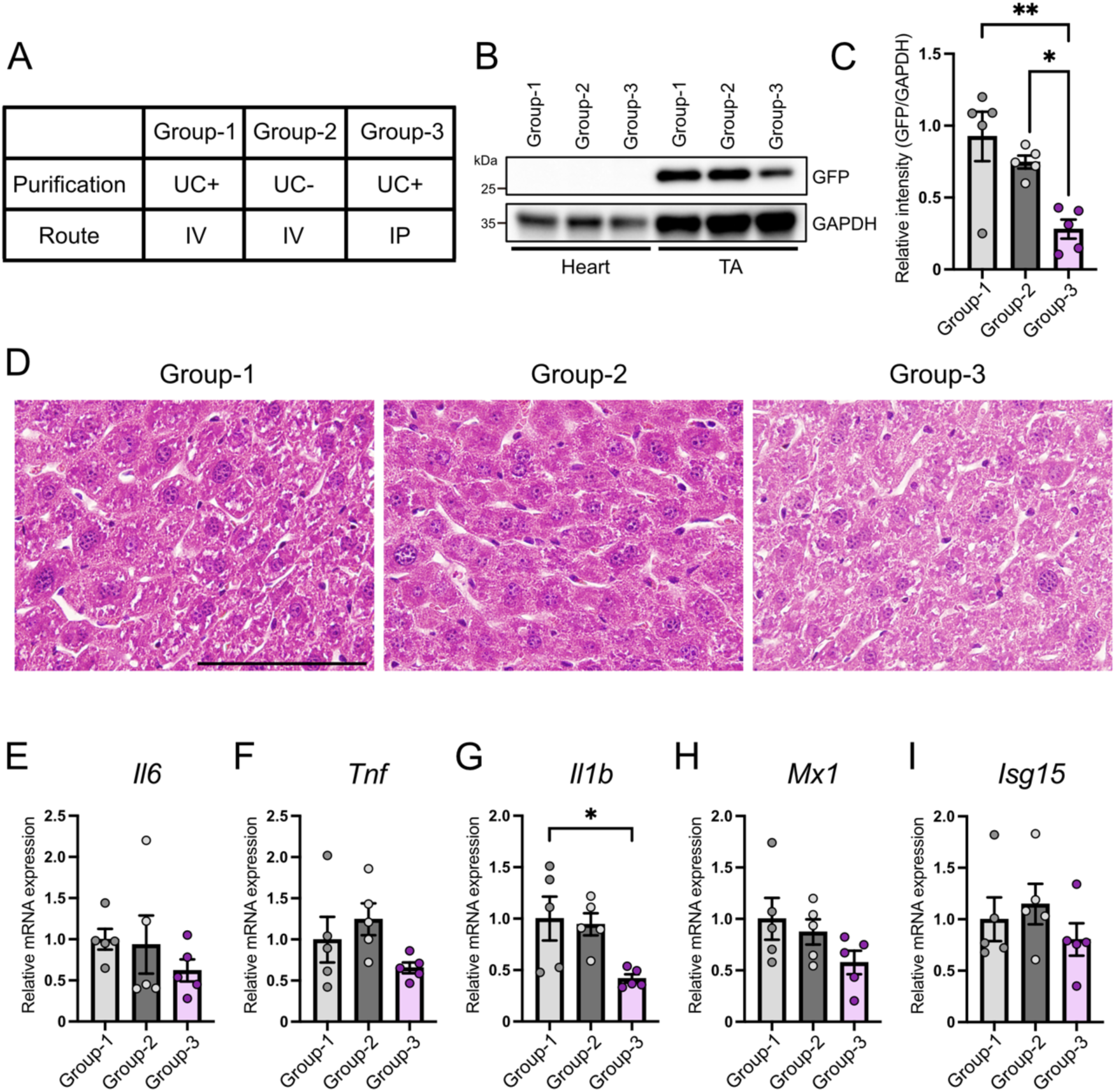
Cardiac detargeting by a miR-208a target site is maintained across purification and delivery workflows. (A) Study design comparing vector purification and injection route. Group 1, UC purified IV; Group 2, kit purified (no UC) IV; Group 3, UC purified IP. (B) Immunoblot analysis of EGFP protein in heart and TA from Groups 1–3. n = 5 mice per group. (C) Densitometric quantification of EGFP protein levels in TA normalized to GAPDH and expressed relative to Group 1 (n = 5). (D) Representative H&E staining of liver sections from Groups 1–3. Scale bar, 100 µm. (E–I) RT-qPCR analysis of inflammatory and interferon response markers in liver: (E) *Il6*, (F) *Tnf*, (G) *Il1b*, (H) *Mx1*, (I) *Isg15* (n = 5). P values were calculated using one-way ANOVA with Tukey’s multiple comparisons test; *P < 0.05, **P < 0.01.

To compare potential tissue responses associated with these workflows, we next evaluated liver histology. Hematoxylin and eosin (H&E) staining of liver sections did not reveal overt differences among Groups 1–3 (Fig. 3D). Consistently, H&E staining of TA sections showed no evident pathological changes across groups (Supplementary Fig. 2A). We further quantified inflammatory and interferon response markers by RT-qPCR in the liver. Expression of *Il6*, *Tnf*, *Mx1*, and *Isg15* did not differ significantly among Groups 1–3 (Fig. 3E, F, H, I). *Il1b* expression was modestly but significantly lower in Group 3 compared with Group 1 (Fig. 3G), whereas no other pairwise comparisons reached significance. In parallel analyses of TA and spleen, the same marker set (*Il6*, *Tnf*, *Il1b*, *Mx1*, *Isg15*) showed no significant differences among groups (Supplementary Fig. 2B–F and Supplementary Fig. 2G–K). Together, these results indicate that the cardiac detargeting strategy implemented in this vector design is robust across purification workflows and dosing routes, whereas skeletal muscle expression is modulated by administration route, with no measurable histological or inflammatory differences among Groups 1–3 under the conditions tested.

### Cardiac detargeting using miR-208a prevents Nrl-induced cardiac hypertrophy and death

To functionally validate this platform in a setting relevant to fast myofiber loss, we focused on large Maf family transcription factors, which serve as key determinants of type IIb myofiber identity in mice and humans (Fig. 4A).^14,25^ Whereas Mafa, Mafb, and Maf have previously been shown to promote the type IIb myofiber program in skeletal muscle, the ability of Nrl to do so has not remained unknown. Nrl retains the conserved transactivation and bZIP domains of large Maf family proteins but is structurally more compact than Mafa, Mafb, and Maf (Fig. 4A). RT-PCR across multiple mouse tissues and skeletal muscles confirmed that Nrl expression is restricted to the retina and undetectable in skeletal muscles and other tissues examined (Fig. 4B).

**Figure 4.**
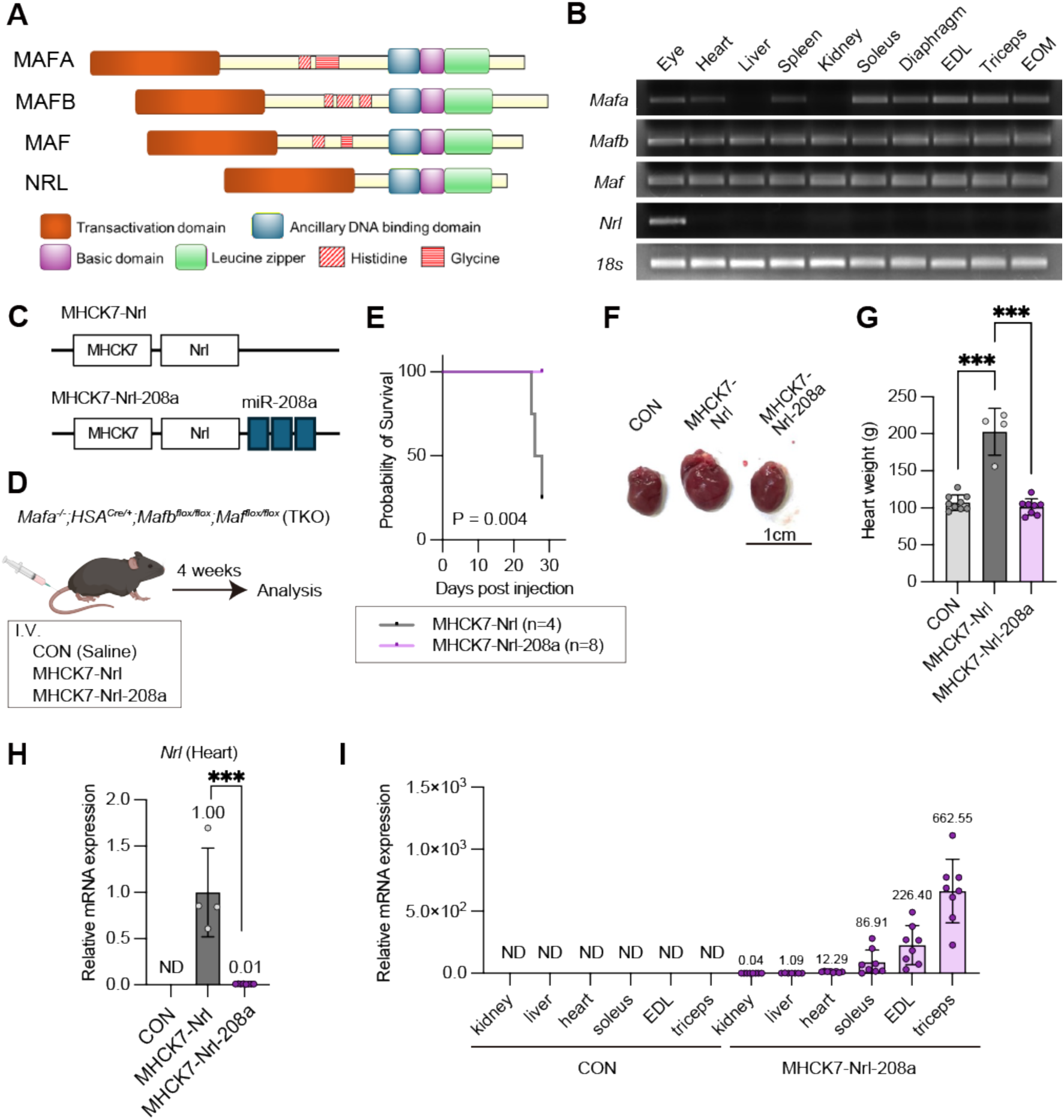
Systemic cardiac-detargeted MyoAAV-Nrl delivery prevents cardiac toxicity and lethality while preserving skeletal muscle expression. (A) Schematic diagram of the protein structure of large Maf family transcription factors. (B) Gel electrophoresis of RT-PCR products corresponding to *Mafa, Mafb, Maf, Nrl*, and *Rn18s* mRNAs across multiple tissues. (C) Schematic of the MHCK7 promoter-driven Nrl cassette packaged in MyoAAV carrying three miR-208a-3p target-site repeats (208a×3) in the 3′UTR. (D) Experimental design for systemic administration of MyoAAV vectors with or without 208a×3 to *Mafa^−/−^; HSA^Cre/+^; Mafb ^flox/flox^; Maf ^flox/flox^* (triple knockout; TKO) mice. (E) Kaplan–Meier survival curves following intravenous injection of MyoAAV-Nrl (n = 4) or MyoAAV-Nrl-miR208a (n = 8). (F) Representative heart images from TKO mice treated with saline (CON), MyoAAV-Nrl, or MyoAAV-Nrl-miR208a. (G) Heart weight in TKO mice treated with CON (n = 9), MyoAAV-Nrl (n = 4), or MyoAAV-Nrl-miR208a (n = 8). (H) RT-qPCR analysis of relative *Nrl* mRNA expression in the heart from TKO mice treated with CON (n = 9), MyoAAV-Nrl (n = 4), or MyoAAV-Nrl-miR208a (n = 8). (I) RT-qPCR analysis of relative *Nrl* mRNA expression in multiple skeletal muscles from TKO mice treated with CON (n = 9) or MyoAAV-Nrl-miR208a (n = 8). P values were calculated using Log-rank (Mantel-Cox) test (E), an unpaired two-tailed t-test (H), and one-way ANOVA with Tukey’s multiple comparisons test (G, I); ***P < 0.001.

Before testing systemic delivery, we first asked whether Nrl possesses intrinsic activity to drive a type IIb myofiber program in vitro and in vivo. To assess the potential relevance of this mechanism to human skeletal muscle, we tested whether ectopic Nrl expression can induce *MYH4* in human skeletal muscle cells. Adenoviral overexpression of NRL in the immortalized human skeletal muscle cell line Hu5KD3^30^ induced *MYH4* expression (Supplementary Fig. 3A), suggesting that Nrl retains the ability to promote a type IIb-associated gene program in human skeletal muscles. Because we previously showed that Mafa, Mafb, and Maf activate *Myh4* transcription through a MAF recognition element (MARE) in the *Myh4* promoter, we next tested whether Nrl could similarly transactivate the *Myh4* promoter using a luciferase reporter containing the predicted MARE sequence. Co-transfection of this reporter with increasing amounts of Nrl expression vector produced a clear dose-dependent increase in normalized luciferase activity compared with empty vector controls (Supplementary Fig. 3B), indicating that Nrl can transactivate the *Myh4* promoter. To further confirm that Nrl is sufficient to induce type IIb myofiber formation in vivo, we first injected MyoAAV-Nrl directly into the soleus muscles of wild-type (WT) mice, in which type IIb myofibers are normally absent. To compare its sufficiency with that of other large Maf family members, we also delivered MyoAAV-Mafa, -Mafb, and -Maf into the soleus muscles and performed immunohistochemical analyses 4 weeks after injection (Supplementary Fig. 3C). Consistent with our previous study, overexpression of Mafa, Mafb, and Maf induced type IIb myofibers in the WT soleus (Supplementary Fig. 3D). Notably, Nrl overexpression produced a similar effect, demonstrating that Nrl is also sufficient to induce a type IIb myofiber program in vivo (Supplementary Fig. 3D). Consistently, *Myh4* expression was robustly upregulated by Mafa, Mafb, Maf, and Nrl overexpression to similar levels (Supplementary Fig. 3E). In contrast, induction of *Actn3*, a direct MARE-dependent target of large Maf factors, was weaker with Nrl than with Maf, although it remained significantly increased relative to control (Supplementary Fig. 3F). *Mybpc2*, another direct MARE-dependent target, was more strongly induced by Mafa and Mafb, whereas Nrl produced a more modest increase (Supplementary Fig. 3G).

Given that Nrl exhibited type IIb-inducing activity in vitro and in vivo, we next examined whether MyoAAV-mediated systemic delivery of Nrl would be feasible. For this purpose, we used large Maf triple-knockout (TKO) mice, in which endogenous Mafa, Mafb, and Maf are depleted in skeletal muscle, as previously described.^14^ Intravenous delivery of conventional MyoAAV-Nrl (Fig. 4C, D) was highly toxic, resulting in death of all treated mice within 1 month (Fig. 4E). Consistent with this severe toxicity, treated mice exhibited marked cardiac enlargement (Fig. 4F, G). Together, these findings suggest that ectopic cardiac expression of Nrl caused severe cardiotoxicity. Because our miR-208a target-site module provides a tunable trade-off between cardiac repression and skeletal muscle expression, we selected three miR-208a target-site repeats for the Nrl vector to maximize the stringency of cardiac detargeting, prioritizing cardiac safety over skeletal muscle expression in this context. We therefore generated an MHCK7 promoter-driven Nrl cassette harboring three tandem miR-208a target sites in the 3′UTR to suppress ectopic cardiac expression (Fig. 4C, D). Notably, incorporation of these miR-208a target sites completely abrogated mortality during the observation period (Fig. 4E) and eliminated detectable cardiac enlargement (Fig. 4F, G). Together, this phenotypic rescue supports the idea that the toxicity observed after delivery of conventional MyoAAV-Nrl was driven, at least in part, by ectopic cardiac Nrl expression. Consistent with these findings, *Nrl* expression was readily detected in the heart after MyoAAV-Nrl administration but was markedly suppressed by the miR-208a target sequence (Fig. 4H). MyoAAV-Nrl-miR208a, however, induced ectopic *Nrl* expression predominantly in skeletal muscle (Fig. 4I). *Nrl* mRNA was undetectable in all tissues examined in uninjected controls. Following injection, only minimal transgene expression was detected in non-muscle tissues such as the kidney and liver, whereas expression in skeletal muscles was markedly higher. Among the skeletal muscle examined, expression was highest in the triceps, followed by the EDL and soleus, revealing a muscle-dependent gradient of transgene expression (Fig. 4I). Together, these results show that miR-208a-mediated cardiac detargeting prevents Nrl-induced cardiac toxicity while preserving efficient delivery to multiple skeletal muscles.

### Cardiac-detargeted MyoAAV-Nrl-miR208a enables systemic induction of the type IIb myofiber program in vivo

Having confirmed by intramuscular injection that Nrl induces a type IIb myofiber program in vivo, we next asked whether this effect could also be achieved by systemic delivery. We intravenously administered MyoAAV-Nrl-miR208 and analyzed myofiber type composition and *Myh4* expression 4 weeks after injection. We quantified myofiber type composition by immunostaining for MyHC isoforms and found that Nrl expression promoted type IIb myofiber formation in multiple muscles, including the EDL, triceps, and soleus (Fig. 5A–F), resulting in a significant shift from IIa/IIx fiber types toward type IIb fibers. This fiber-type shift was accompanied by increased Myh4 mRNA expression (Fig. 5G–I), indicating that systemic delivery of cardiac-detargeted MyoAAV-Nrl-miR208a is sufficient to induce a type IIb-associated transcriptional and myofiber remodeling program in vivo. Together, these findings support Nrl as a functional cargo for counteracting age-associated loss of fast myofibers.

**Figure 5.**
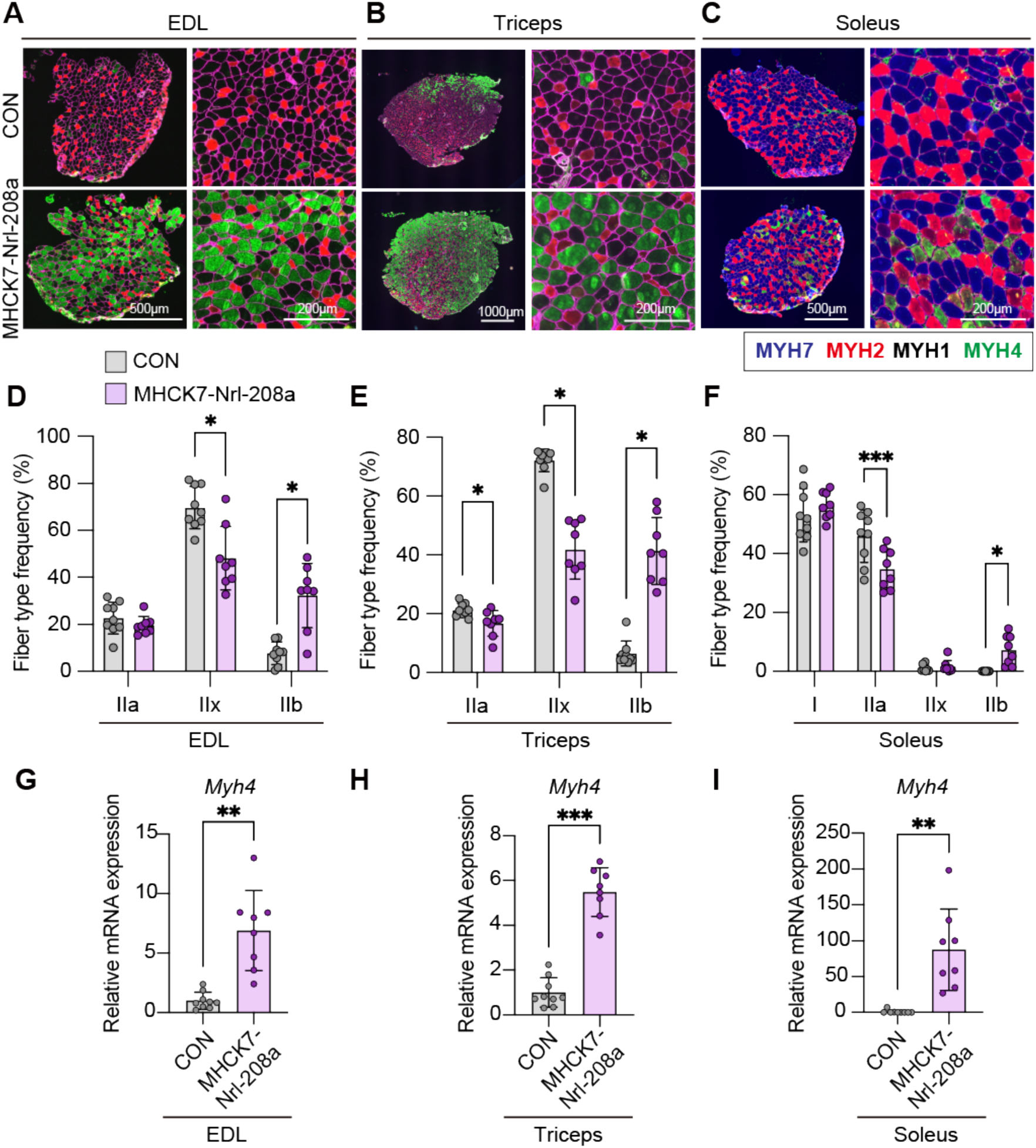
Systemic delivery of cardiac detargeted MyoAAV-Nrl induces type IIb myofibers across multiple skeletal muscles. (A–C) MyHC immunostaining using fiber-type-specific antibodies in the EDL (A), triceps (B), and soleus (C) muscles of TKO mice intravenously injected with saline (CON) or MyoAAV-Nrl-miR208a. Type I fibers are shown in blue, type IIa fibers in red, and type IIb fibers in green. Unstained fibers were classified as type IIx fibers (black). Scale bars are shown in the indicated images. (D–F) The proportion of each fiber type in the EDL (D), triceps (E), and soleus (F) muscles from TKO mice in the CON and MyoAAV-Nrl-miR208a groups (n = 9 and 8, respectively). (G–I) RT-qPCR analysis of relative *Myh4* mRNA expression in the EDL (G), triceps (H), and soleus (I) muscles from TKO mice in the CON and MyoAAV-Nrl-miR208a groups (n = 9 and 8, respectively). P values were calculated using an unpaired two-tailed t-test; *P < 0.05, **P < 0.01, ***P < 0.001.

### Cardiac detargeted MyoAAV-Nrl-miR208a promotes myofiber hypertrophy across multiple skeletal muscles

Because type IIb myofibers generally have a larger CSA than type IIa or IIx myofibers, we next asked whether systemic delivery of MyoAAV-Nrl-miR208a affects not only myofiber type composition but also myofiber size in vivo. Notably, MyoAAV-Nrl-miR208a-treated mice exhibited higher final body weight and greater weight gain than in saline-treated controls (Fig. 6A, B). We then measured myofiber CSA in the EDL, triceps, and soleus muscles. MyoAAV-Nrl-miR208a treatment significantly increased myofiber CSA in all three muscles compared with controls, indicating that Nrl delivery promotes myofiber hypertrophy (Fig. 6C–F). To further determine whether this hypertrophy is linked to Nrl-induced type IIb myofiber remodeling, we analyzed the correlation between the proportion of type IIb myofibers and CSA within each muscle. Interestingly, CSA positively correlated with the proportion of type IIb myofibers in the EDL and triceps, whereas no significant correlation was detected in the soleus (Fig. 6G–I). Together, these results indicate that Nrl-induced type IIb myofiber remodeling is associated with myofiber hypertrophy, particularly in fast muscles such as the EDL and triceps. Collectively, these findings show that systemic delivery of cardiac detargeted MyoAAV-Nrl-miR208a increases body weight and myofiber size, likely through the induction of the fast type IIb myofiber gene program.

**Figure 6.**
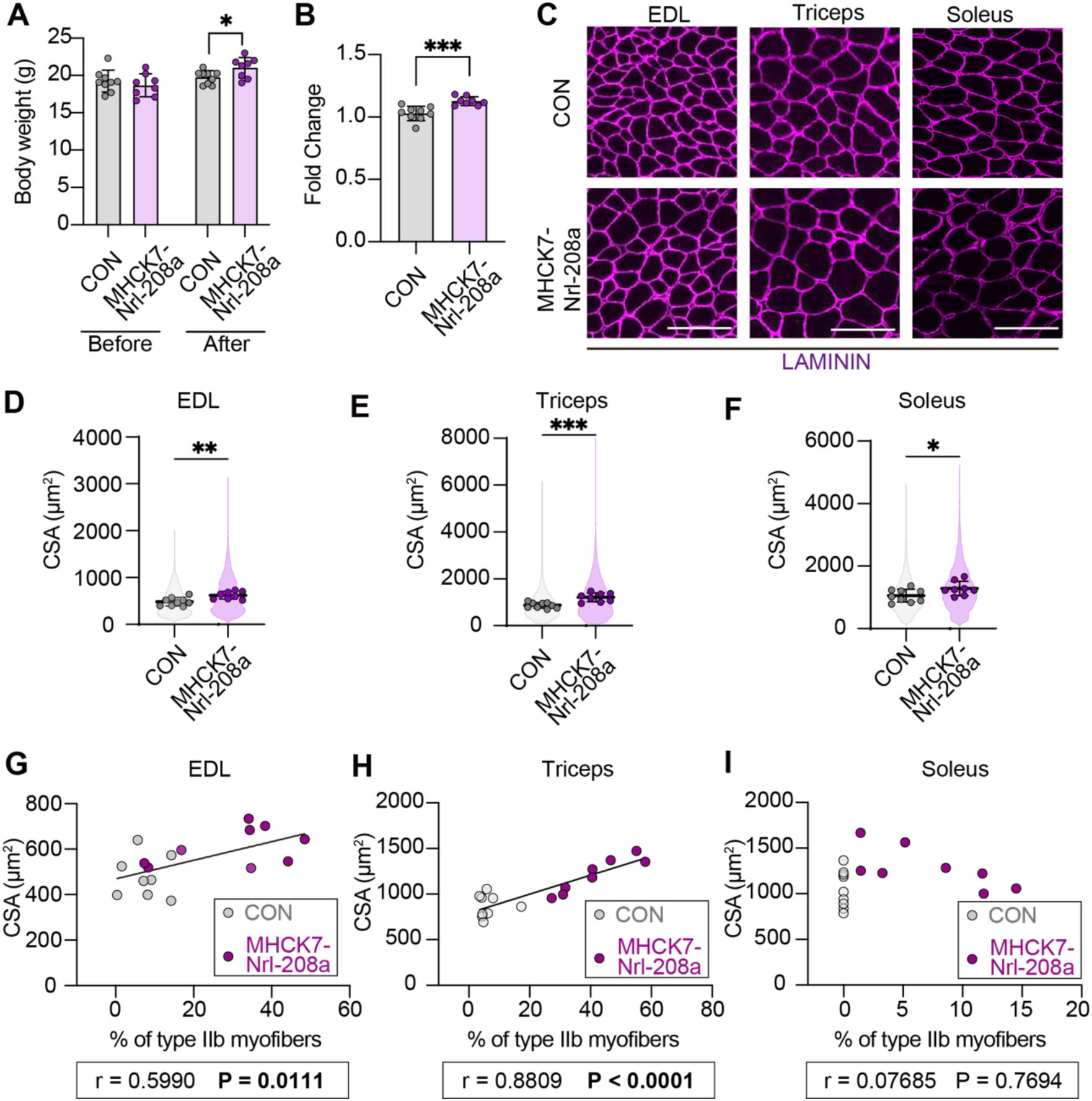
Systemic delivery of cardiac detargeted MyoAAV-Nrl induces myofiber hypertrophy across multiple skeletal muscles. (A) Body weight of TKO mice before and after systemic injection of saline (CON; n = 9) or MyoAAV-Nrl-miR208a (n = 8). (B) The fold change in body weight after systemic injection relative to pre-injection body weight in TKO mice treated with saline (CON; n = 9) or MyoAAV-Nrl-miR208a (n = 8). (C) Laminin immunostaining in the EDL, triceps, and soleus muscles of TKO mice systemically injected with saline (CON) or MyoAAV-Nrl-miR208a. Scale bars, 100 μm. (D–F) Cross-sectional area (CSA) of myofibers in the EDL (D), triceps (E), and soleus (F) muscles from TKO mice in the CON and MyoAAV-Nrl-miR208a groups (n = 9 and 8, respectively). (G) The correlation between CSA and the percentage of type IIb myofibers in the EDL, triceps, and soleus muscles of TKO mice in the CON and MyoAAV-Nrl-miR208a groups. P values were calculated using an unpaired two-tailed t-test; *P < 0.05, **P < 0.01, ***P < 0.001.

## Discussion

MyoAAV is an engineered AAV variant with enhanced muscle tropism, enabling efficient gene delivery to striated muscles including skeletal and cardiac muscle.^13^ These capsids display an RGD-containing peptide on the capsid surface, which enhances integrin-mediated binding and cellular entry and thereby improves systemic skeletal- and cardiac muscle transduction compared with conventional AAV serotypes. However, AAV-based strategies face key constraints, including a limited packaging capacity (≈4.7 kb) that restricts the combined size of regulatory elements and transgene payload. Moreover, achieving skeletal muscle-biased expression remains challenging, because even muscle-active promoters such as MHCK7 can also drive substantial expression in the heart.^31^ Thus, additional strategies are needed to reduce cardiac off-target expression while preserving robust transgene expression in skeletal muscle. In this study, we established a MyoAAV platform incorporating miR-208a target sites to enable robust Nrl delivery to skeletal muscle while suppressing cardiac expression. Systemic administration of the conventional MyoAAV-MHCK7-Nrl construct (lacking miR-208a sites) caused pronounced cardiac transgene expression, cardiac hypertrophy, and high lethality. In contrast, systemic GFP overexpression is well tolerated under our experimental conditions, indicating that observed toxicity is transgene-dependent. These findings underscore the importance of incorporating an additional safeguard against cardiac off-target expression when systemically delivering biologically potent transcriptional factors.

Consistent with previous studies showing that cardiac-enriched miR-208a can be used to reduce unwanted cardiac transgene expression in muscle gene delivery,^32,33^ our results further identify target-site dosage as a key determinant of the trade-off between cardiac suppression and skeletal muscle output in the MyoAAV context. In particular, our comparison of miR-208a×1 and miR-208a×3 showed that target-site dosage critically influences this balance. Although endogenous miR-208a-related myomiRs were detectable in EDL at levels much lower than those in the heart, the miR-208a×3 design reduced transgene expression in skeletal muscle as well as in the heart, indicating that even low-abundance endogenous miRNAs can become functionally significant when multiple target sites are incorporated into the same transcript. Thus, our data support a quantitative model in which the effect of miRNA-mediated detargeting is determined not simply by tissue-specific miRNA presence or absence, but by the balance between endogenous miRNA abundance and target-site dosage, consistent with previous studies on tandem miRNA target-site design.^34–37^

Using this newly developed system, we demonstrated that Nrl, like other large Maf factors, is capable of regulating type IIb myofiber identity in skeletal muscle. In mouse skeletal muscle, Nrl strongly induced *Myh4* expression and type IIb characteristics, with activity nearly comparable to that previously reported for other large Maf factors.^14^ Moreover, the tunable MyoAAV-miR-208a platform enabled efficient systemic delivery of Nrl to multiple skeletal muscle groups while minimizing cardiac expression. Systemic Nrl delivery increased the proportion of type IIb myofibers in both forelimb and hindlimb muscles and was associated with increased body weight and larger myofiber CSA. Collectively, these findings reveal an unanticipated capacity of Nrl to drive a type IIb-like transcriptional program in skeletal muscle and support the utility of the MyoAAV-miR-208a platform as a practical framework for systemic, skeletal muscle-biased gene delivery.

Systemic Nrl overexpression was associated with an overall increase in mean myofiber CSA in the three muscles examined. Because type IIb myofibers are typically the largest myofiber subtype, the expansion of the IIb population could contribute substantially to the increase in bulk CSA. However, we also detected CSA enlargement across multiple fiber types in the triceps and soleus raising the possibility that Nrl influences myofiber hypertrophy through mechanisms that are at least partly independent of IIb induction. Consistent with this idea, large Maf factors have been implicated in muscle phenotypes beyond fiber type regulation: Maf overexpression attenuates denervation-induced atrophy with repression of atrophy-related genes,^23^ and Mafa has been linked to muscle growth in ovine skeletal muscle.^38^

Despite induction of type IIb myofibers and a clear increase in mean CSA following systemic ectopic Nrl expression, whether this remodeling confers functional improvement remains to be elucidated. In our analyses, although Nrl robustly induced *Myh4*, its ability to upregulate other fast contractile components, such as *Actn3* and *Mybpc2*, appeared more limited than that of other large Maf family members. Because skeletal muscle force production depends on coordinated interactions among myosin, actin, and multiple regulatory and structural proteins within the sarcomere,^39,40^ disproportionately elevating a single myosin heavy-chain isoform may be insufficient to enhance whole-muscle performance if other components of the contractile and regulatory machinery are not concomitantly optimized. Consistent with this notion, prior work showed that Actn3 overexpression in mice did not increase muscle strength/performance,^41^ underscoring that augmenting a single fast-associated structural component is not necessarily sufficient to enhance muscle force. Notably, in our in vivo gain-of-function comparisons, Maf most strongly induced *Actn3*, whereas Mafa and Mafb more readily upregulated *Mybpc2*, suggesting that individual large Maf family members may preferentially reinforce distinct modules of the type IIb-associated contractile program.^25^ This raises the possibility that functional reconstruction of a fully “competent” IIb phenotype—structurally and mechanically—may require coordinated engagement of multiple large Maf–dependent transcriptional arms rather than maximal activation of *Myh4* alone. Future studies that systematically tune the expression of individual large Maf factors (or defined combinations) and pair transcriptomic/sarcomeric profiling with direct force measurements will be important to determine whether large Maf engineering can be leveraged not only to shift fiber-type identity and size, but also to improve muscle strength in a predictable functional outcomes.

Importantly, our data also highlight several remaining limitations. Low but detectable transgene transcripts in non-muscle tissues indicate that some off-target expression persists, and we therefore cannot exclude the possibility that such ectopic Nrl expression contributed, at least in part, to systemic phenotypes observed in this study, including increased body weight. In the heart, low but detectable cardiac transgene transcripts remained in the Nrl condition, indicating that residual expression may persist at the mRNA level despite robust suppression at the protein level. Such residual expression may have limited practical consequences for standard transgenes, but becomes more consequential for payloads such as Cre, because even minimal and transient escape from miRNA-mediated repression may be sufficient to trigger irreversible recombination before silencing is fully established. Consistent with this, miR-208a-based detargeting did not prevent cardiac recombination in our Cre reporter assay, even with three target-site repeats, indicating that miR-208a target sites alone are not sufficiently stringent for Cre payloads under our conditions. This outcome is consistent with prior reports showing that the effectiveness of miRNA-mediated detargeting for recombinase payloads can vary substantially depending on vector design and experimental conditions, and that off-target recombination can remain readily observable even when miRNA targeting is implemented.^42^ This context dependence may reflect, at least in part, the underlying biology of the selected miRNA. Because miR-208a is encoded within an intron of the cardiac myosin heavy chain gene *Myh6*, its repressive effect would be expected to be strongest in *Myh6*-expressing cardiomyocytes, potentially allowing limited transcript escape in cardiac cell populations with lower miR-208a activity. Thus, for heart-off, muscle-on Cre applications, achieving sufficient stringency may require detargeting strategies beyond miR-208a and/or additional transcriptional restriction, such as more stringent skeletal muscle-selective promoter designs.

In conclusion, we show that Nrl, the only large Maf family member not detectably expressed in skeletal muscle under physiological conditions, can also promote a type IIb myofiber program. By incorporating miR-208a target sites into a muscle-tropic MyoAAV cassette, we achieved systemic delivery of Nrl with reduced cardiac expression, complete prevention of lethality, and robust induction of type IIb myofibers across forelimb and hindlimb skeletal muscles. These findings expand the functional scope of Nrl beyond the retina and show that cardiac-detargeted MyoAAV-Nrl enables systemic fast myofiber remodeling and hypertrophy across multiple skeletal muscles. Although still speculative, our results also raise the possibility that manipulation of Nrl/large Maf activity could coordinately modulate myofiber composition and muscle mass, with potential relevance to atrophy-prone conditions such as sarcopenia and neurogenic muscle wasting.

## Material and methods

### Mice

Mice were maintained under specific pathogen-free conditions in the Laboratory Animal Resource Center at the University of Tsukuba, Ibaraki, Japan, and the Advanced Medical Research Center for Animal Models of Human Diseases at Fujita Health University, Aichi, Japan, and Meiji University, Kanagawa, Japan. TKO mice were generated in a previous study.^14^ All animal procedures performed at the University of Tsukuba were approved by the University of Tsukuba Animal Ethics Committee (authorization number: 25-157; 25-161), whereas those performed at Fujita Health University and Meiji University were approved by the Institutional Animal Care and Use Committee of Fujita Health University (approval No. APU25060) and Meiji University Animal Experiment Committee (approval No. MUIACUC2022-04). All experiments were conducted in accordance with the relevant institutional guidelines and Japanese regulations. Humane endpoints included severe reduction in spontaneous activity with inability to access food or water and/or signs of respiratory distress, at which point animals were euthanized.

### AAV production

For the EGFP-based experiments, AAV vectors were generated based on the previously described MyoAAV system^13^, in which EGFP expression was driven by the MHCK7 promoter.^28^ To generate cardiac detargeted constructs, either tandem target sequences for miR-208a-3p, miR-208b-3p, and miR-499-5p or one or three copies of the miR-208a-3p target sequence were inserted downstream of the EGFP coding sequence in the 3′ UTR, as indicated in each experiment. AAVs were generated based on the MyoAAV.2A production method described previously.^28^ To compare preparation methods with or without ultracentrifugation, viral particles were purified using the AAVpro Purification Kit Maxi (All Serotypes; Takara Bio) according to the manufacturer’s instructions.

For the Nrl-based experiments, AAVs were generated by triple plasmid transfection of 293T cells using polyethyleneimine (PEI MAX; Polysciences). AAV isolation and purification were performed essentially as previously described.^43^ Titers of AAV vector stocks were determined by quantitative real-time PCR (RT-qPCR) using primers listed in Supplementary Table 1.

### AAV administration for tissue specificity experiments

For tissue specificity experiments, ICR mice purchased from Japan SLC were injected with 1 × 10^12^ genome copies (gc) per animal of the indicated AAV vectors via either intraperitoneal or intravenous administration, depending on the experimental design. In some experiments, AAV preparations purified with or without ultracentrifugation were compared. Four weeks after injection, skeletal muscles, heart, liver, and spleen were collected for analysis. For histological analysis, some mice were perfused with 4% paraformaldehyde (PFA) in PBS before heart collection.

For Cre-dependent reporter experiments, B6.Cg-Gt(ROSA)26Sor<tm9(CAG-tdTomato)Hze>/J mice (Jackson Laboratory, stock no. 007909)^44^, maintained on a C57BL/6N background, were used. Mice were administered approximately 6 × 10^11^ gc of AAV by intraperitoneal injection under isoflurane anesthesia. Fourteen days after injection, hearts were collected and photographed under a fluorescence stereomicroscope.

### Overexpression of large Mafs

For Hu5KD3^30^, the myotubes were transduced with adenovirus at a multiplicity of infection of 250 in differentiation medium for 48 h at two days post-differentiation of Hu5KD3. The medium was then replaced with fresh differentiation medium without adenovirus. The myotubes were used for subsequent experiments at four days post-infection. The adenovirus vectors used to overexpress the large MAF transcription factors in Hu5KD3 cells are as follows: pAV[Exp]-mCherry-CMV > hMAFA, pAV[Exp]-mCherry-CMV > hMAFB, and pAV[Exp]-mCherry-CMV > hMAF, pAV[Exp]-mCherry-CMV>hNRL. pAV[Exp]-mCherry-CMV was used for control. All vectors were constructed and packaged by VectorBuilder Inc. (Chicago, IL, USA), with vector IDs as follows: VB010000-9300kfr (mCherry), VB900139-7528hhe (hMAFA), VB900139-7526uvc (hMAFB), VB900139-7527axe (hMAF), and VB900205-1246kuy (hNRL). The mice were injected with 5.0 × 10^11^ particles (intramuscular injection) / 1.0 × 10^13^ particles (intravenous injection) of MyoAAV. Control mice were injected with the same amount of saline. The muscles were collected four weeks after injection.

### Luciferase assay

A *Myh4* promoter–luciferase reporter construct (−450 to +50) in the promoter-less pGL4.10 vector (Promega) used in this study was previously generated as described.^14^ To examine whether Nrl could activate the *Myh4* promoter, pGL4-Myh4 (100 ng) was co-transfected with pRL-TK vectors (50 ng, Promega) and pRP[Exp]-CMV>mNrl (0 ng, 100 ng, 200 ng) into AAVpro 293T cells (Takara Bio) using the FuGENE HD transfection reagent (Promega). The Nrl expression vector was constructed and packaged by VectorBuilder Inc. (Chicago, IL, USA). The vector ID is VB250807-1222xbu, which can be used to retrieve detailed information about the vector on vectorbuilder.com. Twenty-four hours after transfection, the cells were harvested and assayed using the Dual-Luciferase Reporter Assay System (Promega). Luciferase activity was measured by GloMax® 20/20 Luminometer (Promega) and was normalized to Renilla luciferase activity from the pRL-TK vector.

### Quantitative analysis of transcripts using reverse transcription PCR

For tissue specificity analysis, total RNA was extracted from tissues using the miRNeasy Mini Kit (Qiagen) with DNase treatment. First-strand cDNA was synthesized using PrimeScript RT Reagent Kit (Takara Bio). Quantitative RT-PCR was performed using TB Green Premix Ex Taq II (Takara Bio) on a Thermal Cycler Dice TP800 (Takara Bio). Relative gene expression was calculated using the ΔΔCt method with *Hprt* as the internal control. For miRNA expression analysis, cDNA was synthesized using the miScript II RT Kit (Qiagen), and quantitative PCR was performed using the miScript SYBR Green PCR Kit (Qiagen). *U6* was used as the internal control. For Maf overexpression analysis, total RNA and DNA were extracted from various tissues and skeletal muscles using ISOGEN (NIPPON GENE, Chiyoda, Tokyo, Japan). First-strand cDNA was synthesized using ReverTra Ace® qPCR RT Master Mix (Toyobo). PCR was performed using a THUNDERBIRD Next SYBR qPCR system (Toyobo) and a CFX Opus 384 Real-Time PCR System (Bio-Rad). The relative amount of each transcript was normalized to 18S ribosomal RNA (*Rn18s)* transcripts. The primer sequences are listed in Supplementary Table 1.

### Histological analysis and immunohistochemistry

For GFP immunostaining in tissue specificity experiments, tissues were fixed with 4% PFA in PBS. TA muscles were processed into 10 μm cryosections, whereas hearts were paraffin-embedded and sectioned at a thickness of 3 μm. Sections were stained essentially as previously described.^45^ Briefly, sections were incubated with an anti-GFP polyclonal antibody (1:500, No. 598, Medical & Biological Laboratories (MBL)) followed by the appropriate secondary antibody. For H&E staining in tissue specificity experiments, paraffin sections of TA muscles and livers (3 μm) were stained with hematoxylin and eosin using standard procedures. Images were acquired using a BX51 microscope with a DP70 camera (Olympus) or a MICA microscope (Leica), depending on the experiment.

For muscle cryosection analysis in large Maf overexpression experiments, hindlimb muscles were mounted in tragacanth gum (FUJIFILM Wako Pure Chemical) on cork discs with the Achilles tendon side oriented vertically, rapidly frozen in isopentane cooled by liquid nitrogen, and cryosectioned (8 μm). Immunohistochemistry, fiber-type quantification, and myofiber CSA measurements were performed essentially as previously described.^14^ Briefly, sections were fixed in cold acetone (−20°C), blocked (5% goat serum/1% BSA in PBS with M.O.M. blocking reagent; Vector Laboratories), incubated with primary antibodies overnight at 4°C, and detected with Alexa Fluor–conjugated secondary antibodies (Thermo Fisher Scientific). . Primary antibodies against MYH7 (1:50, BA-D5, DSHB), MYH2 (1:100, SC-71, DSHB), MYH4 (1:100, BF-F3, DSHB), LAMININ (1:200, 4H8-2, Santa Cruz) were used. Images were acquired using a BIOREVO BZ-X800 microscope system and analyzed using the Hybrid Cell Count application (Keyence, Osaka, Japan) to determine fiber-type proportions and CSA.

### Western blotting

For western blot analysis of GFP expression, heart and skeletal muscle tissues from AAV-injected mice were harvested, rapidly frozen, and homogenized in RIPA buffer supplemented with protease inhibitors, essentially as described previously,^45^ with minor modifications. Lysates were clarified by centrifugation, and protein concentrations were measured using a BCA Protein Assay Kit (Takara Bio). Equal amounts of protein were separated by SDS-PAGE and transferred to PVDF membranes. Membranes were incubated with primary antibodies against GFP (1:4000, anti-GFP polyclonal antibody, No. 598, MBL) and GAPDH (1:5000, #2118, Cell Signaling Technology), followed by HRP-conjugated anti-rabbit IgG secondary antibody (1:10000, #7074, Cell Signaling Technology). Signals were detected using ImmunoStar LD (FUJIFILM Wako Pure Chemical) and imaged with a cooled CCD camera system (Light-Capture, ATTO). Band intensities were quantified using ImageJ (version 2.16.0), and GFP expression levels were normalized to GAPDH.

### Quantification and statistical analysis

Graphs are presented as mean ± SEM, as indicated in Figure legends. Significance was calculated using GraphPad Prism software by either Student’s *t*-test or one-way ANOVA with Tukey’s test: *P < 0.05, **P < 0.01, ***P < 0.001.

### Data availability statement

- All unique reagents generated in this study are available from the corresponding author, Ryo Fujita, in accordance with the relevant material transfer agreements.
- This paper does not report original code.
- Any additional information required to reanalyze the data reported in this paper is available from the corresponding author, Ryo Fujita or Keisuke Hitachi upon request.

## Acknowledgments

We would like to thank Erina Fukushima, Chika Ohshima, and the Open Facility Center at Fujita Health University, Japan, for technical assistance. We are grateful to Dr. Tohru Hosoyama (National Center for Geriatrics and Gerontology, Japan) for kindly providing the Hu5/KD3 cell line.

## Funding

This work was supported by the JST FOREST Program (JPMJFR234V to R.F. and JPMJFR225M to K.H.). This work was additionally supported by AMED-CREST (JP23gm171008h to R.F. and S.T.), Grants-in-Aid for Scientific Research (B) (24K02876 to R.F.) and for Challenging Exploratory Research (24K22244 to R.F.) from the Japan Society for the Promotion of Science (JSPS). S.S. was supported by a JSPS Research Fellowship for Young Scientists (23KJ0287). This work was also supported by the Takeda Science Foundation (R.F.).

## Author Contributions

K.H., S.S., and R.F. conceived and designed the study. K.H., S.S., M.W., R.T., A.K., Y.Y., Y.K., and M.I. performed the experiments. K.H., S.S., M.W., R.T., A.K., Y.Y., Y.K., M.I., and R.F. analyzed and interpreted the data. T.K., T.S., S.T., and K.T. provided critical reagents, technical support, and conceptual advice. R.F. and K.H. supervised the project and acquired funding. K.H., S.S., and R.F. wrote the manuscript. All authors reviewed and approved the final manuscript.

## Declaration of Interests

The authors declare no competing interests.

## Notes

### Competing Interest Statement

The authors have declared no competing interest.

