## Supplementary Figures for "Cardiac-Detargeted MyoAAV Enables Systemic Nrl-Mediated Fast Myofiber Remodeling and Hypertrophy Across Multiple Skeletal Muscles"

Keisuke Hitachi *et al*.

Correspondence:
 (Keisuke Hitachi)
 (Ryo Fujita)

**This PDF file includes:**

Figs. S1 to S3

Table S1.

### Supplementary Figure 1

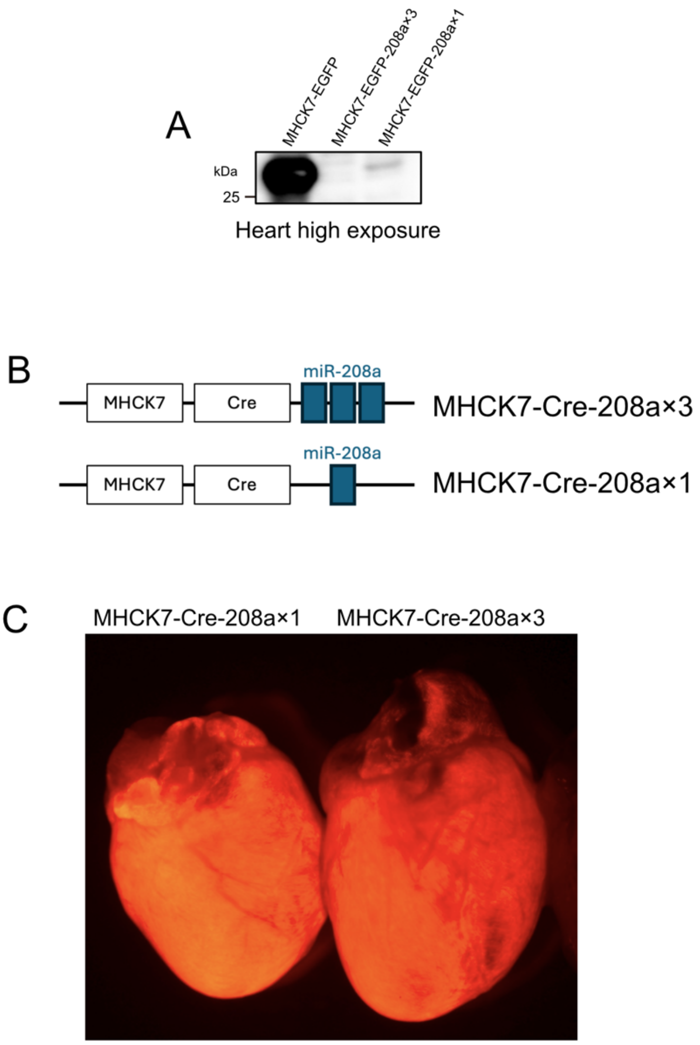


**Supplementary Figure 1. Additional analyses of miR-208a target-site repeat-dependent cardiac detargeting across readouts.**

(A) High-exposure imaging of the EGFP immunoblot shown in Fig. 2E. A faint residual EGFP signal is detectable in the heart in the 208a×1 group under high-exposure conditions. Low- and high-exposure images were acquired from the same membrane. (B) Schematic of the MHCK7 promoter-driven Cre cassette packaged in MyoAAV carrying either three (208a×3) or one (208a×1) miR-208a-3p target-site repeat(s) in the 3′UTR. (C) Representative fluorescence images of hearts from Rosa26-LSL-tdTomato reporter mice after intraperitoneal administration of Cre-208a×3 or Cre-208a×1 vectors, showing tdTomato activation (red) in the heart. n = 3 mice per group.

#### Supplementary Figure 2

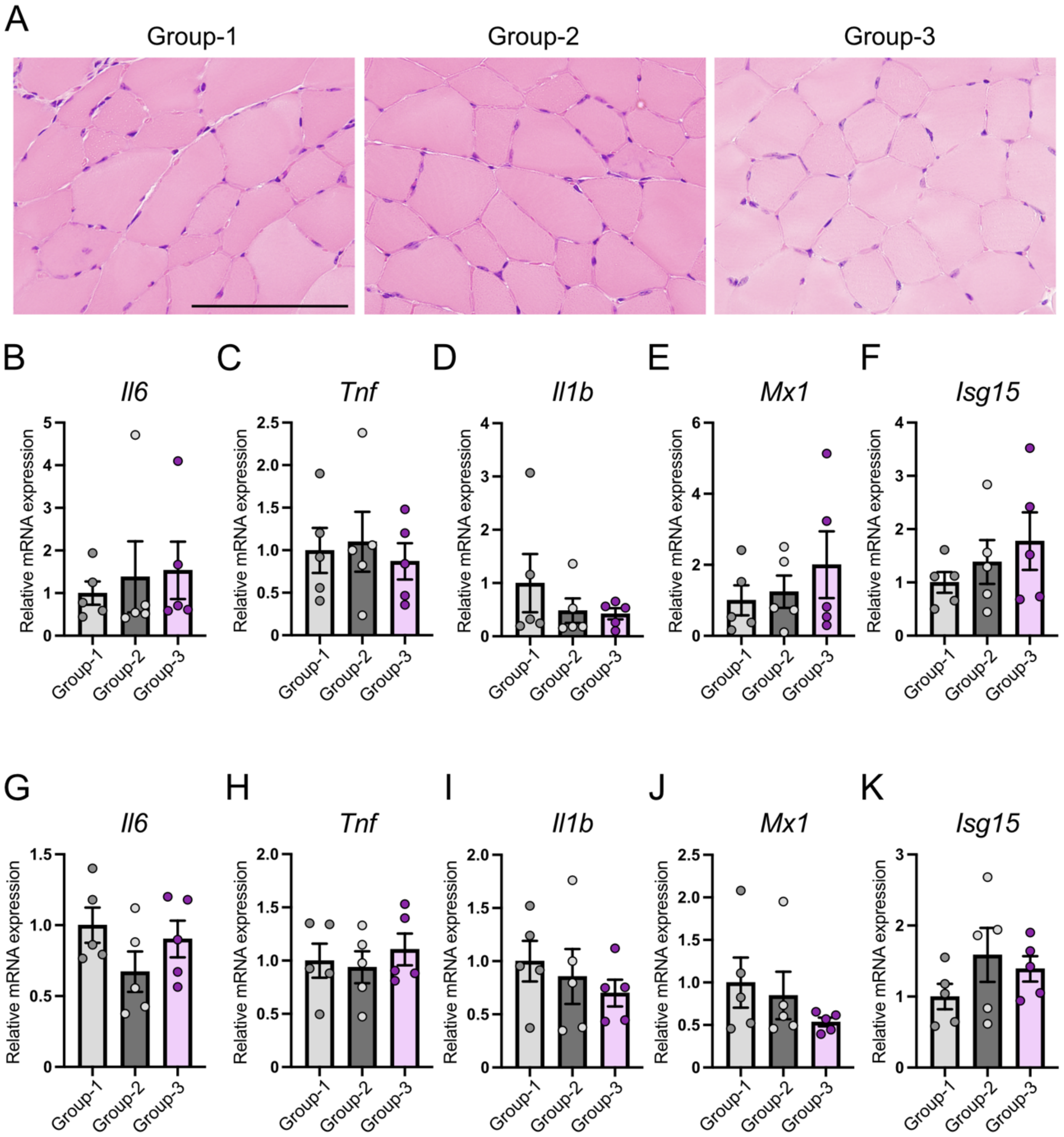


**Supplementary Figure 2. No overt histological or transcriptional inflammatory differences in TA and spleen among the purification and delivery workflows.**

(A) Representative H&E staining of TA sections from Groups 1–3. Scale bar, 100 µm. (B–F) RT-qPCR analysis of *Il6*, *Tnf*, *Il1b*, *Mx1*, and *Isg15* in TA from Groups 1–3. (G–K) RT-qPCR analysis of *Il6*, *Tnf*, *Il1b*, *Mx1*, and *Isg15* in spleen from Groups 1–3. n = 5 mice per group. P values were calculated using one-way ANOVA with Tukey's multiple comparisons test; no statistically significant differences were detected.

#### Supplementary Figure 3

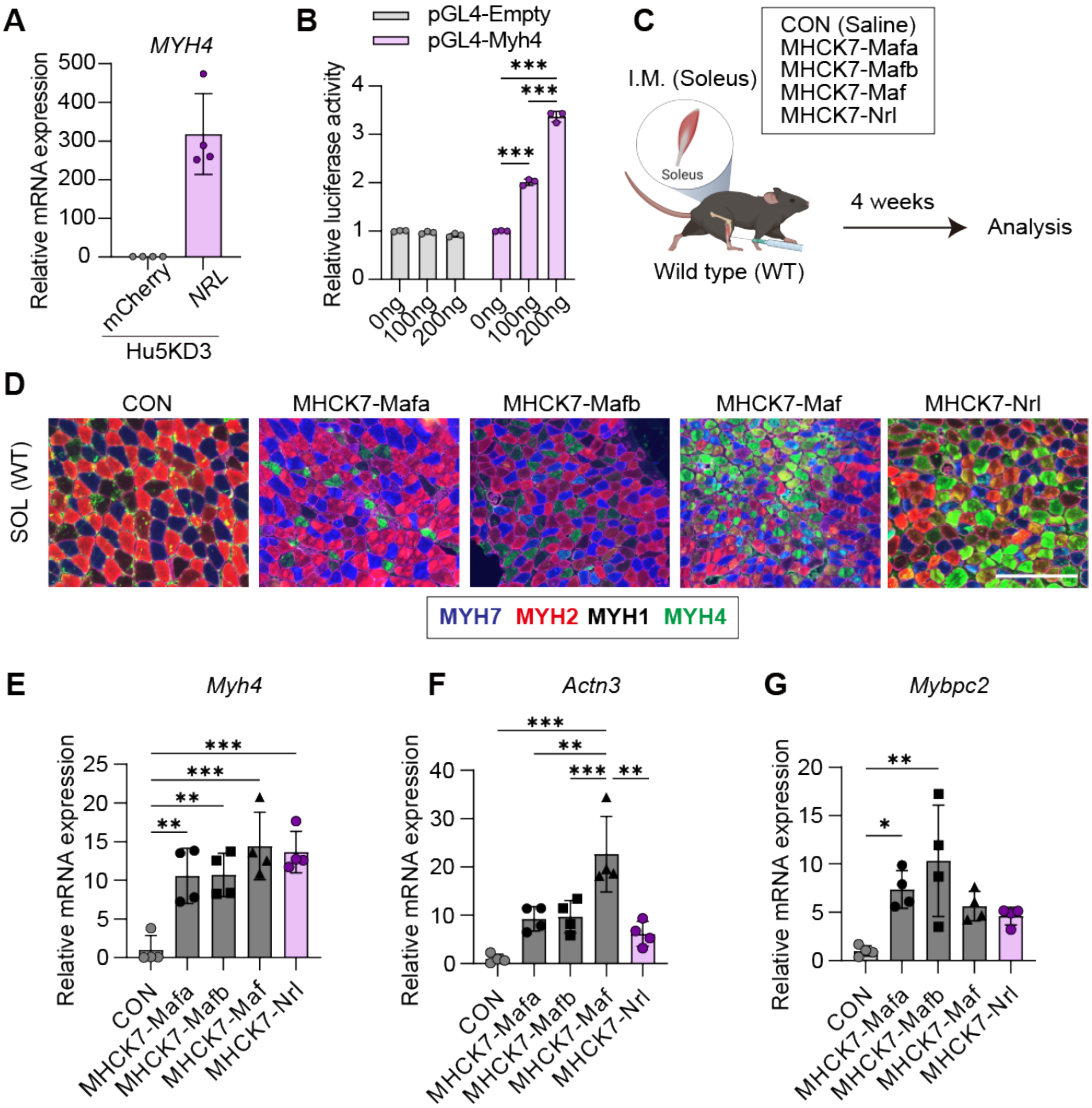


**Supplementary Figure 3. Nrl induces the type IIb myofiber program in vitro and in vivo.** (A) RT-qPCR analysis of relative *Myh4* mRNA expression in Hu5KD3 cells (human muscle cell line) after adenoviral overexpression of mCherry (CON; n = 4) or NRL (n = 4). (B) Relative Myh4 luciferase reporter activity in HEK293T cells transfected with an Nrl expression vector or empty vector (n = 3 per condition). (C) Experimental design for intramuscular administration of MyoAAV-MHCK7-Mafa, -Mafb, -Maf, or -Nrl into wild-type (WT) mice. Saline injection was used as a control (CON). (D) MyHC immunostaining using fiber type-specific antibodies in the soleus muscles of WT mice intramuscularly injected with saline (CON) or MyoAAV-Mafa, -Mafb, -Maf, or -Nrl. Type I fibers are shown in blue, type IIa fibers in red, and type IIb fibers in green. Unstained fibers were classified as type IIx fibers (black). Scale bars, 200 μm. (E–G) RT-qPCR analysis of relative *Myh4* (E), *Actn3* (F), and *Mybpc2* (G) mRNA expression in the soleus muscles from WT mice in the CON, MyoAAV-Mafa, MyoAAV-Mafb, MyoAAV-Maf, and MyoAAV-Nrl groups (n = 4 per group). P values were calculated using one-way ANOVA with Tukey's multiple comparisons test or an unpaired two-tailed t-test; *P < 0.05, **P < 0.01, ***P < 0.001.

**Table S1. List of primer sequences**

| Gene | Fw | Rv |
|---|---|---|
| *miR-208a-3p* | ATAAGACGAGCAAAAAGCTTGT | |
| *miR-208b-3p* | ATAAGACGAACAAAAGGTTTGT | |
| *miR-499-5p* | TTAAGACTTGCAGTGATGTTT | |
| *U6* | miScript Primer Assay (Qiagen, MS00014000) | |
| *Il-6* | TAGTCCTTCCTACCCCAATTTCC | TTGGTCCTTAGCCACTCCTTC |
| *Tnf* | CAGGCGGTGCCTATGTCTC | CGATCACCCCGAAGTTCAGTAG |
| *Il1b* | GAAATGCCACCTTTTGACAGTG | TGGATGCTCTCATCAGGACAG |
| *Mx1* | GACCATAGGGGTCTTGACCAA | AGACTTGCTCTTTCTGAAAAGCC |
| *Isg15* | GGTGTCCGTGACTAACTCCAT | TGGAAAGGGTAAGACCGTCCT |
| ITR | GGAACCCCTAGTGATGGAGTT | CGGCCTCAGTGAGCGA |
| *Mafa* | AGGCCACCACGTGCGCTTGG | GCTGCTGCACCCGCTTGAAG |
| *Mafb* | CAACGGTAGTGTGGAGGACC | CTTCTGCTTCAGGCGGATCA |
| *Maf* | GCAATGAACAATTCCGACCT | CCGGTTCCTTTTTCACTTCA |
| *Nrl* | TCACCCACCTTCAGTGAGC | CCCGAGAACCTCATCCGAC |
| *Myh4* | CACCTGGACGATGCTCTCAGA | GCTCTTGCTCGGCCACTCT |
| *Actn3* | GAGAAACAGCAGCGGAAAAC | GAAATGACCTCCAGGAGCAG |
| *Mybpc2* | ATTCGTAGGTGACCGAGTGG | CGTCCTTCTTGAAGCGGTAG |
| *18s* | AGTCCCTGCCCTTTGTACACA | GATCCGAGGGCCTCACTAAAC |
| *Hprt* | TCAGTCAACGGGGGACATAAA | GGGGCTGTACTGCTTAACCAG |
| *MYH4 (human)* | AAAGGTGGCCATTTACAAGCT | CAGCAGAGTTCAGACTTGTCAG |
| *TBP (human)* | TGTATCCACAGTGAATCTTGGTTG | GGTTCGTGGCTCTCTTATCCTC |
